# Investigating the Conformational Flexibility of Staphylokinase Across Multiple Time Scales

**DOI:** 10.64898/2026.02.03.703606

**Authors:** Anthony Legrand, Linda Kasiarova, Naina Verma, Jan Mican, Joan Planas-Iglesias, Pavel Kohout, Alan Strunga, Josef Kucera, Tomas Henek, Pavel Vanacek, Tim de Martines, Pavel Kadeřávek, Lukáš Žídek, Jiří Damborský, Stanislav Mazurenko, David Bednar, Lenka Hernychova, Zbyněk Prokop, Martin Marek

## Abstract

Cardiovascular diseases, including ischemic stroke, necessitate improved thrombolytic agents. A microbe-encoded plasminogen activator staphylokinase (SAK) is a promising alternative to the widely used tissue plasminogen activator (tPA) due to its high fibrin specificity and low production cost. To overcome potential immunogenicity hampering its use in clinical settings, the low-immunogenic variants SAK SY155 and SAK THR174 were previously engineered. However, the molecular basis underlying their reduced immunogenicity is not understood and requires detailed elucidation. Here, we determine molecular structures and compare flexibility between low-immunogenic and immunogenic SAK variants, using a combination of experimental and computational structural techniques. Our analyses show that all variants share the canonical SAK fold and retain similar plasminogen activation kinetics, despite the number of introduced substitutions. Crucially, the low-immunogenic variants exhibit distinct flexibility profiles, with SAK THR174 showing substantially increased flexibility in the H1 helix and B3 region. SAK SY155 exhibits an increased flexibility in the H1-B3 loop and propensity to homodimerize. These flexibility changes are found in the known immunogenic epitopes. Our multi-scale flexibility analysis provides the molecular explanation for the reduced immunogenicity, altered thermostability, and retained fibrinolytic function of the engineered variants. This information is critical for the design of next-generation thrombolytics.

## Introduction

Cardiovascular and cerebrovascular diseases remain the leading causes of mortality in the world, with stroke alone accounting for approximately 6.5 million deaths annually^1^. As the global burden of stroke is projected to rise, particularly in developing countries^2^, there is an urgent need for more effective and accessible stroke treatments. Ischemic stroke, the most common form, results from a disruption in cerebral flow, caused by occlusion of vessels by fibrin-rich thrombi^3^. While the current standard treatment, relying on recombinant tissue plasminogen activator (rtPA, alteplase), can restore perfusion^4,5^, it is limited by a short half-life, poor fibrin specificity, high production cost, and a significant risk of bleeding^6,7^. Staphylokinase (SAK), a bacterial plasminogen activator, offers a promising alternative to alteplase. It selectively activates fibrin-bound plasminogen via complex formation with already active plasmin, offering better fibrin specificity and potentially reduced systemic side effects. Its small size and low-cost recombinant production make it especially attractive for use in resource-limited settings^8,9^. However, its clinical application is hindered by strong immunogenicity in humans^10,11^.

To overcome this limitation, several SAK variants have been engineered to reduce immune recognition while preserving fibrinolytic activity. One such variant, SAK SY155 (SY), was developed through rational design based on B-cell epitope mapping and incorporates twelve substitutions aimed at reducing antibody binding^12^. Another example, SAK THR174 (THR), developed by ThromboGenics and later commercialized by Rhein Minapharm Biogenetics, was similarly designed to lower immunogenicity^13,14^. Previous studies have identified immunogenic “hotspot” residues – K35, E38, E80, D82, K74, E75, R77^15,16^ and A72-E75, N95-E99, W66, K135, E19, K102, K121^17^, suggesting that helix H1 and β-strands B3 and B5 form the core of the SAK epitope^12^. However, delineating the structural basis underlying immunogenicity remains a challenge.

Although the wild-type SAK (WT) has been characterized by X-ray crystallography^18^ and NMR^19^, structural data on the SY and THR variants are lacking. Moreover, there is limited information about the flexibility of any of these proteins. Only one co-crystal structure of a SAK-microplasmin complex has been determined^20^, leaving substantial gaps in our understanding of SAK action and how engineered mutations affect its conformational behaviour and function.

In a previous study, we dissected the thrombolytic mechanism through kinetic measurements and mathematical modelling, showing that SAK must first bind to plasmin (Plm) to form an active heterodimer that subsequently converts additional plasminogen (Plg) into Plm, thereby potentiating fibrinolysis^8^. In this work, we determine molecular structures and compare flexibility profiles between the WT and two low-immunogenic variants SY and THR, using X-ray crystallography, NMR spectroscopy, hydrogen-deuterium exchange mass spectrometry (HDX-MS), and molecular dynamics (MD) simulations. These results offer new insights into the relationship between sequence, structure, flexibility, and immunogenicity in engineered SAK variants.

## Material and methods

### Gene synthesis and molecular cloning

SAK STAR WT, SAK STAR SY155 and SAK STAR THR174 genes, hereafter referenced to WT, SY and THR, respectively, were assembled by *de novo* synthesis (GeneArt ThermoFisher,USA), streamlined for transcription, and adapted to *E. coli* codon usage. Gene synthesis also included the addition of flanking restriction sites required for convenient subcloning into the expression vector pET21b. The vector bears the T7 promoter, lac operator, ampicillin resistance gene for selection, and a lacI repressor for tight regulation of gene expression. It also includes an N-terminal hexahistidine tag for the affinity purification of the recombinantly expressed proteins followed by TEV protease cleavage sequence.

### Protein overproduction and purification

*E. coli* strain BL21(DE3) containing recombinant plasmid pET21b-N-His-WT/SY/THR was grown in 10 mL Lysogeny broth (LB) complemented with 100 mg/mL Ampicillin (Amp) overnight at 37°C, 180 rpm. The preculture was inoculated into 1 L LB with the same concentration of Amp, cultivated at 37°C until reaching an optical density 0.6 (at 600 nm), at which point production was induced with 0.5 mM IPTG. Production was carried out for 4 hours at 37°C, 120 rpm, and cells were harvested by centrifugation at 4000 g, 4°C, for 30 min. The pellet was resuspended in 40 mL purification buffer A (K_2_HPO_4_, KH_2_HPO_4_, NaCl, Imidazole 10mM, pH 7.5) and frozen at -70°C. The frozen culture was then thawed in water, 80 µL of DNase (1mg/mL) was added to the culture, and the culture was sonicated with three 2-minute cycles with an amplitude 50% and pulse on/off time 5s. The disintegrated culture was centrifuged at 20 000 g, 4°C for 1h. The supernatant was filtered using a syringe filter of 0.45 µm and the protein was purified by affinity chromatography (AKTA Pure, Cytiva). Protein was eluted into purification buffer B (K_2_HPO_4_, KH_2_PO_4_, NaCl, Imidazole 500 mM, pH 7.5) Proteins were then dialyzed into PBS buffer (Na_2_HPO_4_, KH_2_PO_4_, KCl, NaCl, pH 7.4) overnight with TEV protease (1.5 mg TEV/100mg protein) for the cleavage of the N-terminal His-tag. Affinity purification was performed again, and this time the flow-through was collected. (AKTA Pure, Cytiva). The collected flow-through was then subjected to size-exclusion chromatography on a 75 16/600 HiLoad Sephadex column (AKTA Pure, Cytiva), and pure protein was collected.

For NMR samples, cells were grown in 10 mL LB over day at 37°C, then transferred to 100 mL M9 minimal salt medium overnight (for 1 L of M9 culture medium: 12.8 g Na_2_HPO_4_ 3 g KH_2_PO_4_ 0.5 g NaCl 1 mM MgSO_4_ 100 µM CaCl_2_ 10 µM ZnCl_2_ 1 µM FeCl_2_ 50 mg/L ampicillin 10 mL, 100× MEM vitamin mix (Sigma-Aldrich M6895) 2 g D-glucose-^13^C_6_ (Cortecnet CC860P10) 1 g 15NH_4_Cl (CN80P10). The main culture was started in 1 L (for WT) or 2 L (for SY and THR) M9. The rest of the expression protocol and the purification were performed as described above.

### Protein quality controls

Purity was checked by 10% acrylamide Tris-Glycine SDS-PAGE. Concentration was determined by absorbance at 280 nm, assuming molar extinction coefficients of 18,910 M^- 1^.cm^-1^ for WT and SY, or 20,400 M^-1^.cm^-1^ for THR (Expasy ProtParam). Protein hydrodynamic radii were measured at 37°C by FIDA, in a standard 1 m (length) / 75 µm (inner diameter) uncoated capillary, at 400 mPa. Protein thermal stability was assessed using nanoDifferential Scanning Fluorimetry (nanoDSF) on a Prometheus instrument at 1°C/min. Melting temperatures (Tm) were determined using the first derivative of the 350/330 nm fluorescence ratio curve. CD spectra were obtained at 0.2 mg/mL final concentration of WT, SY and THR from 195 to 280 nm at 1nm bandwidth with 0.5 s integration time as an average of 3 readings using a Chirascan spectropolarimeter (Applied Photophysics, UK).

### NMR structure determination

Experiments were carried out at the Josef Dadok National NMR Centre, at CEITEC, Brno, Czechia. WT structure was analysed on a 950 MHz (^1^H Larmor frequency) Bruker Avance NEO NMR spectrometer with a 5 mm triple resonance (^1^H/^19^F-^13^C-^15^N) cryoprobe and cooled ^1^H and ^13^C preamplifiers. SY155 and THR174 structures were analysed on a 700 MHz (^1^H Larmor frequency) Bruker Avance NEO NMR spectrometer with a 5 mm triple resonance (^1^H-^13^C-^15^N) cryoprobe, and cooled ^1^H, ^13^C, and ^15^N preamplifiers. All pulse programs are from Bruker’s catalogue. Temperature was 25°C. Structure determination was based on NMR measurements performed on ^15^N- and ^13^C-labelled protein (at 600 uM for WT, 466 uM for SY, or 439 uM for THR) in 50 mM sodium phosphate 100 mM NaCl 1mM DTT 0.1 mM EDTA pH = 6.7 adjusted with 10% D_2_O. Relevant acquisition and processing parameters are provided in **Table S2**.

Spectra were converted from Bruker to UCSF format using bruk2ucsf from Poky^21^. Semi-automated backbone assignment was performed using Poky, I-PINE, and NMRtist^21,22^, and lists of backbone chemical shift statistics were generated Then, NMRtist was employed for final peak picking, chemical shift assignment, and structure determination. In the case of WT, additional upper limit distance restraints (UPLs) were generated and refined using Poky and AUDANA (on Poky servers) then used as additional input for UPL refinement using NMRtist (structure determination application). The final structure ensemble was analysed with, and Ramachandran plots were plotted, using Molprobity^23^. Folded regions were labelled by majority rule (at least 10 states with the same label out of 20 states) from DSSP annotation of each state^24^.

Structure alignment and visualisation was done with PyMol Molecular Graphics System (version 3.0.4 Schrrlldinger, LLC) using: *intra_fit & resi 24-134* for WT and resi 25-135 for SY155 and THR174 (from the first to the last residue in a secondary structure element, according to DSSP annotation). This gives an RMSD for every state against the first state (i.e. the state of lowest energy), the average of which being defined as the “RMSD vs best state”. “RMSD vs mean” was defined as:

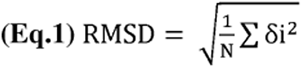

where N is the number of states (N=20) and δi is the distance to average position, thus representing the RMSD of every state against a hypothetical “mean” structure.. For comparison, SY and THR were aligned to WT using: *align {structure}, 2sak & resi 25-134*.

### NMR relaxation measurements

Experiments were carried out at the Josef Dadok National NMR Centre, at CEITEC, Brno, Czechia, on a 600 MHz (^1^H Larmor frequency) Bruker Avance NEO NMR spectrometer with a 5 mm quadruple resonance (^1^H-^31^P-^13^C-^15^N) inverse cryoprobe and cooled ^1^H and ^13^C preamplifiers. Pulse programs are described in^25^ and use heat compensation (so the same amount of irradiation is applied during each experiment). Temperature was adjusted to 25.0°C using 99.8% MeOD as an external standard^26^.

Relaxation experiments were performed on ^15^N-labelled protein (at 300 uM for WT, 428 uM for SY, or 210 uM for THR), in 50 mM sodium phosphate 100 mM NaCl 1mM DTT 0.1 mM EDTA pH = 6.7 adjusted with 10% D_2_O. Relevant acquisition parameters are provided in **Table S2**. The following relaxation delays were used for R1 measurement: 20, 260, 60, 660, 260, 500, 100, 340, 160, 260, and 860 ms and for R2 measurement: 14.4, 129.6, 57.6, 72, 100.8, 43.2, 86.4, 72, 28.8, 115.2, and 72 ms. For heteronuclear steady-state NOE measurements we applied ^1^H saturation achieved with π pulses of duration 200 µs and Q3.1000 shape^27^, with a 2294.6 Hz offset from water to match their frequency to the center of the amide proton region, and far off-resonance (45 kHz) for heat compensation during the measurement of reference spectra. hetNOE, R1, and R2 datasets were processed using NMRPipe: data conversion was done using the *bruker* module, and spectral processing used a classical 2D processing script. We used the hetNOE analysis module from Poky’s Notepad. R1 and R2 peak heights were fitted with a monoexponential decay function to compute R1 and R2 parameters using SciPy Python library. Errors come from random noise sampling from the hetNOE analysis module, or from standard fitting error and propagated error from one triplicate (260 ms for R1 and 72 ms for R2 data). Reduced spectral density mapping (RSDM) was performed as described previously^28^. Errors on spectral densities J(ω) were propagated from hetNOE, R1, and R2 errors using standard algebra.

### Protein crystallization

Initial crystallization screening was performed in 96-well protein crystallization plates (UVXPO 2 Lens Crystallization Plate, SWISSCI) using a Gryphon crystallization robot (Art Robbins Instruments). 0.1 and 0.2 µL of pure protein solutions were mixed with 0.1 µL of the reservoir solution in ratios 1:1 and 1:2 and equilibrated against 100 µL of a reservoir solution in a sitting-drop vapor diffusion arrangement. Nine commercial crystallization screens, namely JSCG-plus, SG1, WIZARD, PACT premier, MEMGOLD Eco, Morpheus MD1-46, Morpheus III, MIDAS (Molecular Dimensions), and INDEX (Hampton Research) were tested. Plates were stored in Crystallization Incubator Rumed Type 3001 (Molecular Dimensions Ltd) at 18°C. The growth of crystals was monitored first after 48 h then weekly using an optical microscope Olympus SZX16 model SZX2-ILLK (Olympus, Japan) equipped with Olympus SDF Plapo 1xPF and 2xPF objectives and Canon EOS 1100D camera (Canon, Taiwan). Pictures were processed in QuickPhoto Camera 3.2 software. Conditions identified in the high-throughput screening to provide crystallization hits were further optimized and scaled up to improve the size and quality of crystals. Final crystals of both SY and THR were grown in a hanging drop arrangement in 36% PEG3350 and 0.18 M sodium citrate (pH 4.4).

### X-ray data collection and structure refinement

All data were collected at 100K temperature at MASSIF-3 beamline at ESRF in Grenoble (France) at wavelength 0.967 Å using DECTRIS EIGER x 4M detector. The X-ray frames were processed and scaled using XDS^29^ and merged using Aimless^30^. The contents of the asymmetric unit were estimated by calculating Matthews coefficient^31^. The structures of SY and THR were solved by molecular replacement with Phenix^32^ software. For the SY variant, an AlphaFold-predicted model^33^ was used as a search model. For the THR, a newly solved X-ray structure of SY was used as a search model. The initial models were refined through several cycles of manual building using Coot^34^ and automated refinement with Phenix^32^. The final models were validated using tools provided by Coot. Ligands were built using eLBOW^35^. Structural panels in figures were generated with PyMOL Molecular Graphics System (Version 3.0 Schrodinger, LLC). Atomic coordinates were deposited in the Protein Data Bank (www.pdb.org) under the PDB IDs 9IAU for SY and 9IAV for THR.

### HDX-MS

The HDX-MS analysis of WT, SY, and THR was carried out in apo-state. A phosphate buffer composed of 10 mM Tris-HCl, 50 mM NaCl, and pH 7.4 was used to prepare the non-deuterated samples. The same composition of D_2_O buffer with pD 7.4, pH 7.0 was used to prepare the deuterated samples. The deuterated samples were labeled for 30, 300, 668, 900, and 1002 s with D_2_O. Each time point was prepared by diluting the protein stock to 2 μM by adding 55 μL of H_2_O/D_2_O buffer. The exchange reaction was quenched by adding 55 μl of 1 M glycine with pH 2.3, further diluting the sample to 1 μM. Fully deuterated control was prepared for each protein sample to correct for the back exchange using the procedure described elsewhere^36^. All samples were prepared using the Leap HDX robotic station (Trajan Scientific and Medica) in technical triplicate.

For each quenched sample, 100 μL was directly injected into an immobilized dual protease column, composed of *Aspergillus niger* prolyl endoprotease and pepsin (AnPep/Pep, Affipro) in 1:2 at a flow rate of 200 µL/min with the loading buffer (0.1% formic acid). The peptides were then trapped and desalted on-line on a peptide microtrap (Phenomenex UHPLC Fully porous polar C18, 2.1 mm) for 3 min before elution using the microtrap (Phenomenex UHPLC Fully porous polar C18, 2.1 mm) for 3 min before elution onto an analytical column (Phenomenex Luna Omega Polar C18 analytical column 1.6 µm, 100 x 1.0 mm, 100 Å), maintained at 2°C. A 6-min linear-gradient (10%-45%) with 80% acetonitrile with 0.1% formic acid was used to separate the peptides (Agilent 1290) at 50 uL/min. The eluate from the analytical column was directed into the TOF mass spectrometer (Bruker Daltonics) with an electrospray ionisation. Blank injections were performed between each sample injection to prevent the carryover of peptides between runs. Mapping samples were analysed in MS/MS mode with trapped ion mobility, while the HDX samples were analysed in MS mode without trapped ion mobility.

LCMS data were first analysed using Data Analysis version 6.1 (Bruker Daltonics) for peak picking and then exported. Tandem mass spectra were searched using MASCOT against the cRAP and contaminants protein database (http://ftp.thegpm.org/fasta/cRAP), containing the sequences of SAK variants, with precursor ion mass tolerance set at 10 ppm and fragment ion mass tolerance at 0.05 Da. No enzyme specificity was applied, with a maximum allowance of two missed cleavages, and no fixed or variable modifications. The false discovery rate at the peptide identification level was set to 1%. A sequence coverage map was prepared using DigDig software, version 0.8.1^37^, and peptide redundancy was calculated. The MS raw files of deuterated and non-deuterated samples, along with the list of peptides and their parameters (score, retention time, charge, sequence, and mobility), were analyzed using HDExaminer version 3.4.1 (Sierra Analytics) (Konermann *et al*., 2011). The software analyzed peptide-level deuteration over different time points and calculated the score based on the fit of theoretical and actual isotope clusters. Confidence levels were assigned as high, medium, or low. Peptides were manually reviewed to improve the fit, and only high- and medium-confidence peptides were included in further calculations. PyHDX v0.4.3^38^ was then used to calculate the amino acid- level of deuteration as relative fractional uptake (RFU), and differential RFU, ΔRFU, allowing comparison of deuterium uptake across states. RFU was calculated by weighted averaging, where each residue’s RFU was determined by averaging RFUs of overlapping peptides, weighted by the inverse peptide length, and visualized in a linear bar plot. The mass spectrometry HDX data have been deposited to the Proteome Xchange (PX) Consortium^39^ via Proteomics Identifications (PRIDE)^40^ partner repository with the dataset identifier PXD072585; Username; Password: eLOcJZIiMDd9.

### MD simulations

The three-dimensional structure of SAK variants WT, SY, and THR, were modelled using the AlphaFold3 server^33^ using default settings. Titratable residues were protonated and orientations of Asn, Gln, and His side chains were optimized using the H^++^ server at 0.1 mM salinity and pH 7.4, with the rest of the parameters left at default values^41^. After processing with H^++^, coordinated crystallographic waters were added to the structure from the structure of the WT ternary complex with microplasmin^18^ (PDB ID: 1BUI) determined using X-ray crystallography at 2.65 Å resolution.

Input topologies and coordinates were prepared using the Tleap module of AMBER 22^42^. The protein parts of the system were parametrized using the ff19SB force field^43^. The system was solvated in a box of water molecules so that all protein atoms were at least 9 Å from the box’s surface so the system is contained within the periodic box. The OPC water model was used^44^. The systems were neutralized using either Na^+^ or Cl^-^ ions so the system has a charge of 0 for the first round of equilibration. The number of ions added for the production simulations was determined using the average volume in the last stage of equilibration simulation to achieve a final salinity of 0.1 M.

Energy minimizations and MD simulations were performed by using the PMEMD.CUDA module of AMBER16^42^. Initially, the system was minimized by 10 steps of steepest descent followed by 9990 steps of conjugate gradient with 500 kcal×mol^-1^.Å^-2^ restraints on all atoms of the protein. The system was further minimized in four more rounds, each consisting of 2500 steps of steepest descent and 7500 steps of conjugate gradient minimization with decreasing harmonic restraints. The restraints were applied as follows: 500, 125, and 25 kcal.mol^-1^Å^-2^ on all backbone atoms of the protein. Finally, the system was minimized with 5000 steps of steepest descent and 15000 steps of conjugate gradient minimization without any restraints. The subsequent MD simulations employed periodic boundary conditions, the Particle Mesh Ewald method to treat electrostatic interactions with long-range cutoff of 10 Å, a 10 Å cut-off for nonbonded interactions, the SHAKE algorithm to fix all bonds containing hydrogens, and a 2 fs time step^45^. Equilibration simulations consisted of two steps: (I) 500 ps of gradual heating from 0 to 310 K using the Langevin thermostat with a collision frequency of 5.0 ps^-1^ and constant volume, (II) 400 ps in the NPT ensemble using the Berendsen barostat at constant pressure of 1.0 bar and pressure relaxation time of 1.0 ps^-1^ and harmonic restraints of 150.0 kcal×mol^-1^Å^-2^ on the positions of all protein backbone atoms^46–48^. Then, the system was further equilibrated during 8800 ps at 310 K in the NPT ensemble as in step (II) in 10 rounds of 400 ps each with decreasing harmonic restraints. The restraints were applied as follows: 100.0, 75.0, 50.0, 25, 15.0, 10.0, 5.0, 1.0, 0.5, and 0.0 kcal×mol^-1^Å^-2^ on the backbone atoms of the protein. After equilibration, 200 ns long production MD simulations were run using the same settings as the last equilibration step. Coordinates were saved at intervals of 2 ps. The simulations were run in 3 replicas, producing 600 ns of aggregated simulation time per each SAK variant. Production trajectories of the MD simulations were analysed using the CPPTRAJ ^49^ module of AMBER14. Trajectories were reimaged from periodic boundary conditions, aligned to the first frame, and used to calculate B-factors of backbone heavy atoms, all using residues 25-136, representing the ordered part of SAK. The simulations were visually analysed using PyMOL Molecular Graphics System (Version 3.1.3. Schrrlldinger, LLC).

### Adaptive sampling MD simulations

SAK variants for which a crystal structure was available were modelled based on the crystal. SAK was sourced from crystal structures (2SAK for WT, 9IAU for SY155, and 9IAV for THR, respectively). Missing N-terminal residues were filled in using an AlphaFold model^33^ of the protein trimmed to the missing residues. Structures were merged to the terminus using PyMol Molecular Graphics System (Version 3.1.3. Schrrlldinger, LLC) and used as templates for final models, modelled using Swiss Model^50^. The structures were thereafter processed using the High Throughput Molecular Dynamics package (HTMD v3)^51^. For each variant, the system was protonated using the systemPrepare module at a pH of 7.8 before solvating in a box of TIP3P water^52^ with the protein at least 10 [ from the box edges using the solvate module. The model was then built using the amber.build module with the ff14SB^53^ force field in AMBER with Na^1^ and Cl^-^ ions added to a concentration of 0.1 M. The system was minimized first using 500 steps of conjugate gradient. Thereafter, the system was heated to 310K and minimized for 2,500,000 steps (10 ns) of NPT equilibration with the Langevin thermostat with a 4 fs time step with results written every 0.1 ns. Of this minimization time, 1,250,000 steps (5 ns) were performed with 1 kcal×mol^-1^×Å^-2^ constraints on the α-carbon and 0.1 kcal×mol^-1^×Å^-2^ constraints on all other atoms. The remaining 1 250 000 steps (5 ns) of minimization were performed with the same settings, but without any constraints. Hydrogen atom mass was scaled by factor 4 and the simulations employed particle mesh Ewald method for treatment of interactions beyond 9 Å cut-off. 50 ns production simulations were then run for 10 epochs with 10 runs per epoch using RMSD of the α-carbon to the input structure as the sampling metric. B-factors were generated using CPPTRAJ from the AMBER16 module to determine per-residue B-factors and RMSD. Simulations were checked for excess RMSD indicating erroneous output.

### Flexibility prediction by Flexpert

The sequences of WT, SY, and THR were used as input to run the inference algorithm of Flexpert v1 ^54^ Sequential module^55^. Flexpert is an ML-based tool trained on the ATLAS MD dataset and provides a per-residue flexibility prediction, ranging between 0 and 1, based on its training on molecular dynamics simulation data.

### Conformational ensembles prediction by BioEmu

For each of the four systems, we generated conformations using BioEmu^56^ a machine-learning-based ensemble generator, employing Model 1.0 from the official GitHub repository (https://github.com/microsoft/bioemu). To remove physically implausible structures, we applied the BioEmu filtering script, which excludes conformations based on Cα–Cα and C–N distances between sequential residues and identifies atomic clashes between atoms of different residues. After filtering, 300 conformations per system were retained for subsequent analysis. RMSF were calculated for Cα atoms after alignment to a representative reference structure (lowest average pairwise RMSD to all ensemble conformations); alignment was performed using Cα atoms while excluding the first and last five residues to minimize terminal flexibility effects, whereas RMSF were computed for all residues.

### Flexibility data projection to 3D structures

Different flexibility values were considered for comparison: i) the directly obtained from crystallography (see section X-ray data collection and structure refinement); ii) B-factors calculated from the Root Mean Square Fluctuations (RMSFs) from the NMR ensembles (section NMR structure determination) according to the following formula:

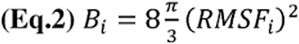

where B_i_ is the resulting B-factor for the i^th^ amino acid represented in the system, and RMSF_i_ is the RMSF for that amino acid calculated through the ensemble of solved conformations; iii) J(0) as a proxy for rigidity also obtained from NMR (section NMR relaxation measurements); iv) relative fractional uptake values obtained from HDX-MS (section HDX-MS); v) B-factors calculated from RMSFs obtained from classic Molecular Dynamics simulations (section MD simulations); vi) Adaptive sampling MD (ASMD) simulations (section Adaptive sampling MD simulations); vii) predicted values from Flexpert (section Flexibility prediction by Flexpert); and viii) B-factors calculated from RMSFs obtained from ML-based predicted conformational ensembles from BioEmu (section Conformational ensembles prediction by BioEmu).

In order to make the different collected flexibility values comparable among systems and methods, all values obtained by any of the aforementioned techniques were internally standardized per system. Thus, each value from a SAK variant prediction series was transformed in the following equation:

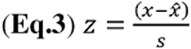

where z is the normalized per-residue value, x is the predicted value (e.g., Flexpert score), x is the average of predicted values and s is their standard deviation.

### Correlation studies

The flexibility collected data, normalised as indicated in the previous section, was concatenated in two different ways: i) methods (crystallographic values, NMR B-factors,NMR J0s, HDS-MS, MD, ASMD, Flexpert, and BioEmu) were concatenated to obtain three different series corresponding to each of the SAK variants, or ii) flexibility values obtained by a single method for the different proteins (WT, SY, THR) were concatenated to obtain eight different series corresponding to each of the methods used to measure or predict flexibility. These series were analyzed with the R package corrplot^57,58^ using Spearman correlation after removing columns with missing data. The subset of residues for which NMR relaxation experiments had trouble obtaining reliable data was analyzed independently using the same method.

## Results

### Low-immunogenic SAK variants share the canonical fold

We successfully achieved mass production and purification of WT (∼ 20 mg per litre of LB medium), SY, and THR (∼ 10 mg per litre of LB medium). WT and THR are strictly monomeric, while SY tends to oligomerise (**Figure 1A**). Monomeric fractions were isolated. CD spectroscopy indicated identical secondary structure content between all isoforms (**Figure 1B**). Fitting of flow-induced dispersion analysis (FIDA) taylorgrams yielded single hydrodynamic radius values: 2.38 nm for WT, 2.31 nm for SY, and 2.56 nm for THR. This indicates that WT and SY are monomeric, while THR could be more prone to dimerisation. The presence of spikes from half the elution time until the elution time reveals the presence of SY and THR aggregates (**Figure 1C**). Similarly, SY and THR are less thermostable than WT by roughly 20°C (**Figure 1D**). We acquired high-resolution crystal structures of SY and THR and compared them to the previously reported WT structure (**Table 1**).

**Figure 1.**
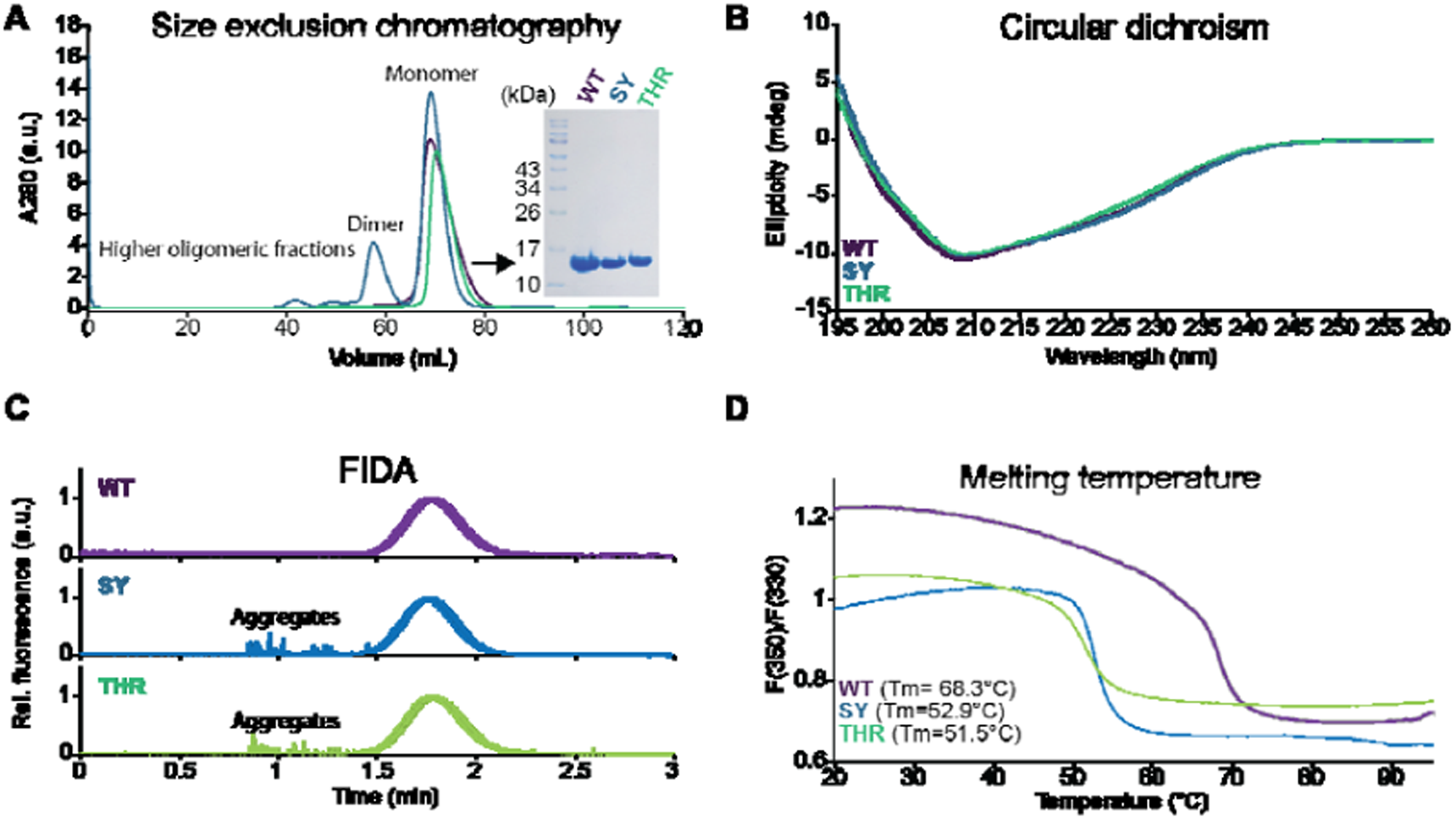
Low-immunogenic variants are less monomeric and thermoresistant. (A) SEC profiles on a HiLoad 16/600 Superdex 200 pg at 12°C. Inlet: 10% Tris-Glycine SDS-PAGE. (B) CD spectra of 13 µM SAK at 25°C. (C) FIDA taylorgrams of 5 µM SAK at 37°C and 400 mPa. (D) Fluorescence ratio at 350 nm over 330 nm of 1 g/L (64 µM) SAK during 1°C/min temperature increase. WT=68.3°C, SY=52.9°C, THR=51.5°C. Colour coding: violet for WT, blue for SY, and green-cyan for THR.

**Table 1.** List of X-ray and NMR structures determined and/or used in this study.

| Protein | Technique | Resolution (Å) | PDB (BMRB) | Reference |
| --- | --- | --- | --- | --- |
| WT | X-ray | 1.8 | 2SAK | Rabijns *et al.*, 1997 |
| WT | NMR | 0.4 | 9RKK (34999) | This study |
| SY | X-ray | 2.1 | 9IAU | This study |
| SY | NMR | 1.1 | 9RB5 (34996) | This study |
| THR | X-ray | 2.5 | 9IAV | This study |
| THR | NMR | 1.2 | 9RKL (35000) | This study |

Overall, all three proteins share the same SAK fold (**Figure 2A**). Notable differences with SY and THR are a split in β-strand B2 into B2 and B2’, and a longer helix H1 (**Figure 2B**). Matthew’s coefficients for SY and THR indicated the presence of four and two molecules per asymmetric unit, respectively, and these were identified in molecular replacement searches. In the case of SY, four chains are organised into two homodimers. PISA analysis^59^ of SY indicated that each dimer interface involves one α-helix and one β-strand from each monomer, with most interface residues located in the α-helix (**Figure 2C**). The SY variant forms dimers both in the crystal (**Figure 2C**) and in solution (**Figure 1A)**. PISA analysis of THR identified a putative dimerization interface dominated by β-strand interactions (**Figure 2D**). However, we did not detect any THR dimer in solution by SEC analysis (**Figure 1A**). In addition, both SY and THR show increased aggregation propensity relative to the WT protein (**Figure 1B**). The C_α_ of P42 is displaced in mutants compared to WT, with different phi and psi dihedral angles (−43.5° and 112.2° in WT, -21.6° and 142.7° in SY, and -72.7° and 140.7° in THR), thus deforming the β-strand B2 (**Figure 2B,E,F**). WT helix H1 is irregular: its C-terminal turn is narrower, indicating that this helix is flexible and might undergo bending motion. However, this feature is absent in the SY and THR variants. We hypothesized that, while structural differences between low-immunogenic and immunogenic SAK proteins seen in crystallographic structures are minimal, they could differ in terms of flexibility profile. Thus, we also determined NMR structure bundles for WT, SY, and THR (**Table 1**). Structural comparison between crystal and NMR structures reveals a striking difference in helix flexibility: the H1 helix in WT is not straight, but rather bends towards the protein’s core, whereas the corresponding helices in SY and THR do not. The bending of the WT helix observed here is consistent with a previously published NMR structure of WT^19^. Moreover, the NMR RMSD of the helix is poorer for the helix and the H1-B3 loop compared to neighbouring residues before H1 (**Figure 3C**), hinting at an increase in available conformational space, and thus an increase in protein flexibility at the helix (**Figure 2G-I**).

**Figure 2.**
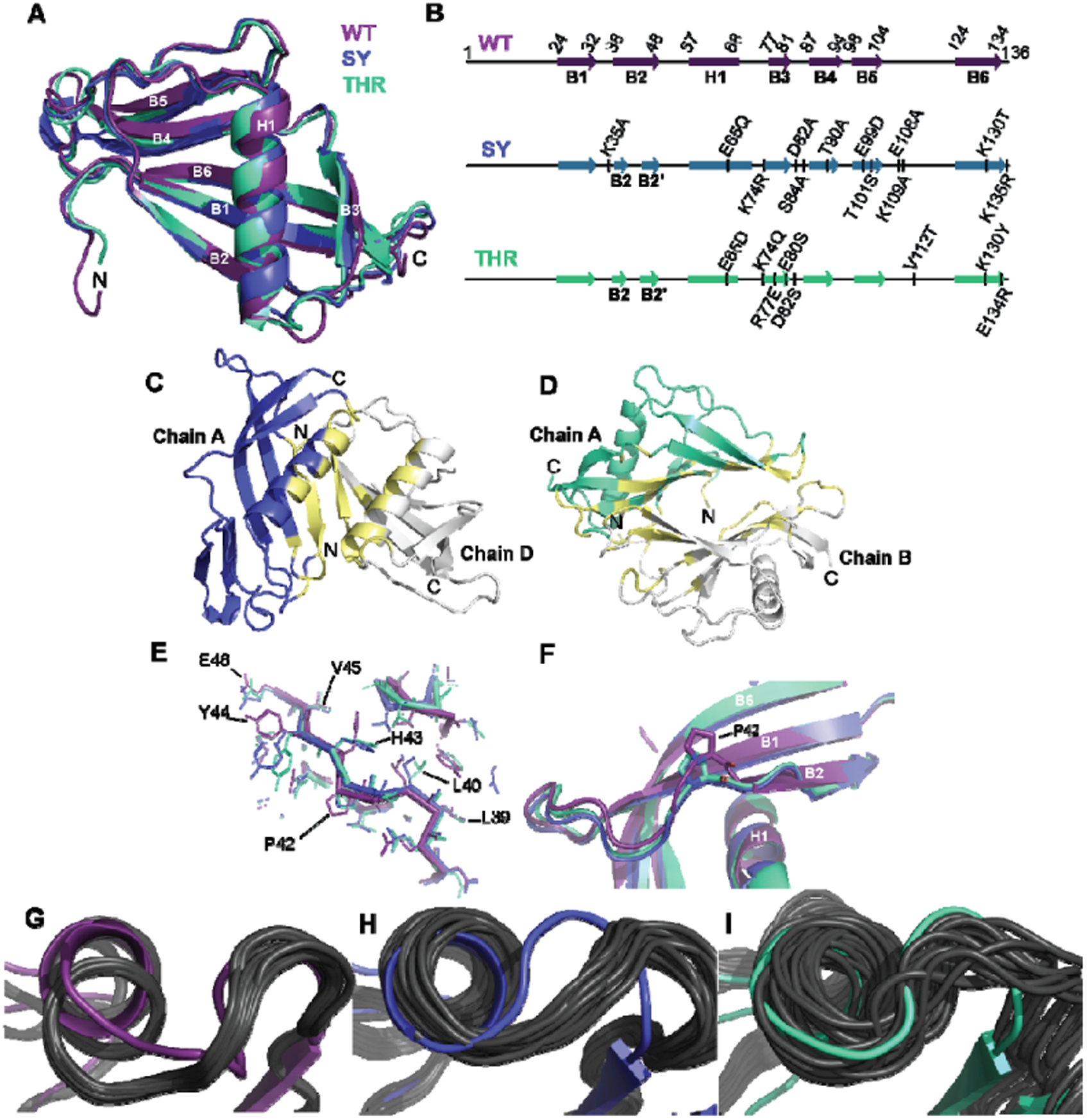
SAK WT and low-immunogenic variants share the same fold. (A) Overview of crystal structures (2SAK,9IAU,9IAV). (B) Secondary structure assignment plots from crystal structures generated with ESPript^60^ (C-D). Crystal dimers of SY (C) and THR (D). Yellow: PISA annotation of residues involved in putative dimerisation interface. (E-F) Comparison of strand B2 between isoforms. Sticks represent backbone, lines represent side chains. (G-I) Cartoon view of helix H1 from its C-terminal side to its N-terminal side. Superimposition of crystal (colour) and NMR structures (black) for WT (F), SY (G), and THR (H). Violet: WT, blue: SY, THR: green-cyan.

**Figure 3.**
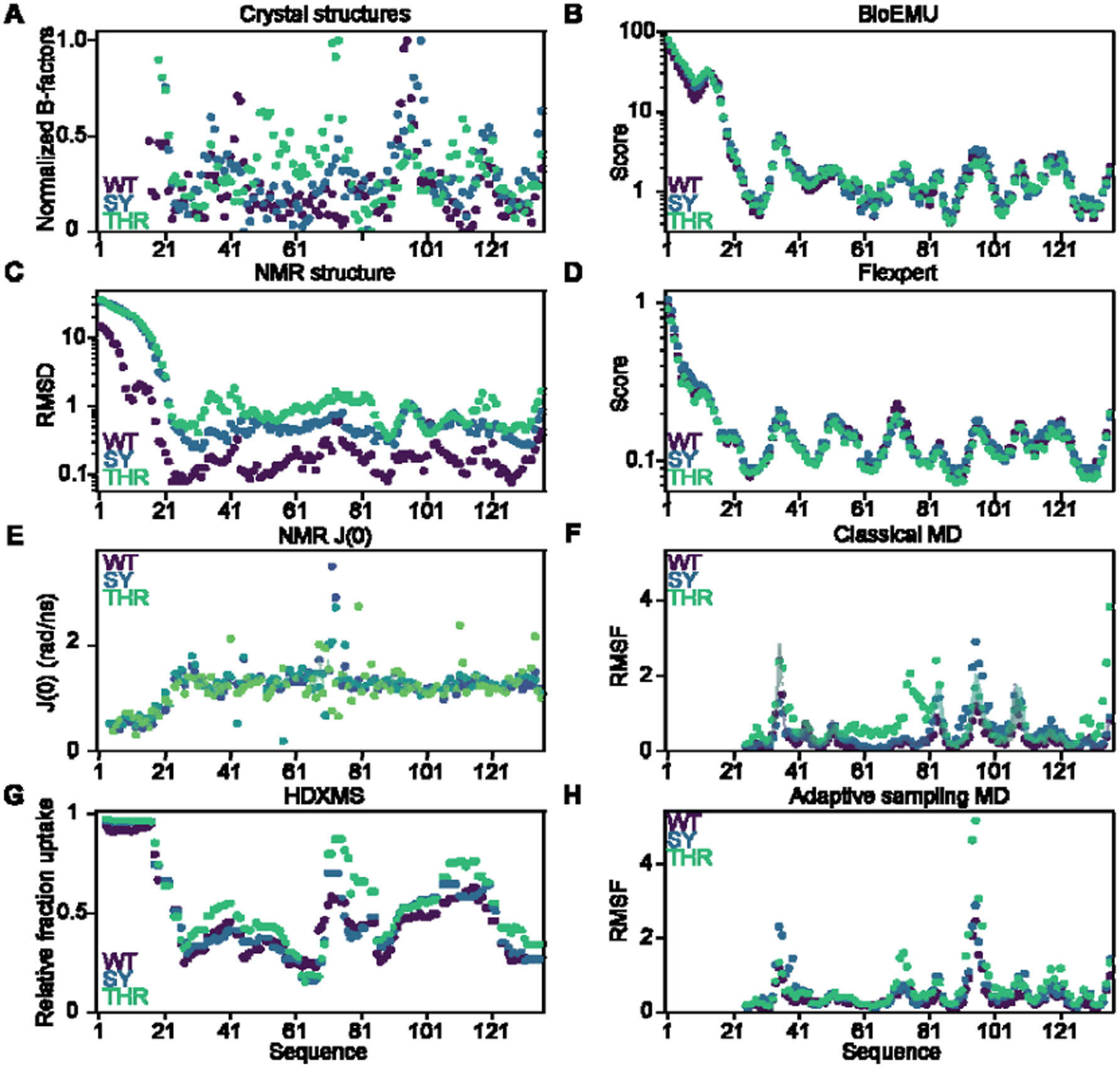
Flexibility measurements reveal different flexibility profiles between SAK variants. (A) B-factors from X-ray crystal structures. (B) BioEMU ensemble-based flexibility prediction. (C) RMSD from NMR structure bundles. (D) Flexpert flexibility prediction. (E) Spectral density J at 0 Hz from NMR relaxation (note that, in contrast to other measurements, it is a measure of protein rigidity). (F) RMSF and standard deviation thereof from classical MD simulations carried out in triplicate. (G) Relative fractional uptake of deuterium from HDXMS. (H) RMSF from adaptive sampling MD simulations. For both classical and adaptive MD simulations, frames were aligned between residues 25 and 136.

### Low-immunogenic variants exhibit different protein flexibility

To further investigate SAK flexibility, we employed a variety of tools: (i) B-factors from crystal structures (fs-ps, very fast motion) (**Figure 3A**), (ii) BioEMU^56^ conformational space sampling (generative deep learning model of protein equilibrium ensembles) (**Figure 3B**), (iii) RMSD of NMR structure bundles (sampling of the conformational space satisfying NMR distance and dihedral restraints) (**Figure 3C**), (iv) Flexpert prediction (ML-based protein flexibility predictor) (**Figure 3D**), (v) NMR spectral density map at 0 Hz J(0) from relaxation measurements (a measure of protein rigidity at the ns timescale, fast motion) (**Figure 3E**), (vi) classical molecular dynamics (MD) simulations in triplicates (ns, fast motion) (**Figure 3F**), (vii) hydrogen-deuterium exchange mass spectrometry (HDX-MS) (s-min, slow motion) (**Figure 3G**), and (viii) adaptive sampling molecular dynamics (ns-µs, fast to intermediate motion) (**Figure 3H**).

All techniques report a highly flexible N-terminal intrinsically disordered domain (IDD) from residues 1 to 24. For X-ray crystallography (**Figure 3A**) and MD simulations (**Figure 3F,H**), IDDs are invisible or excluded from structure alignment and RMSF calculations, respectively. NMR RMSD of WT IDD is slightly lower than that observed in SY and THR (**Figure 3C**). There are very few long-range (more than four residues apart) NOESY distance constraints in the IDD: 3.63 constraints per residue in WT, 1.67 constraints per residue in SY, 1.92 constraints per residue in THR. The higher number of long-range distance constraints in WT IDD could explain its lower RMSD and is a common bias of using NMR RMSD to estimate protein flexibility: other factors, such as peak overlap and relaxation, can influence the number of distance restraints too. Hydrogen-deuterium relative fractional uptake (RFU) is close to 1 for residues 1 to 17, then decreases sharply until residue 27, indicating extensive solvent access and a lack of stable secondary structure where backbone protein-protein H-bonds would form and inhibit exchange with deuterium (**Figure 3G**). NMR J(0), a measure of protein rigidity at the ns timescale, is 2-3 times lower in the IDD compared to the globular domain (0.4-0.5 rad/ns vs 1-1.5 rad/ns) (**Figure 3E**), which starts at residue 25. In contrast, crystal B-factors are very noisy, where the only clearly identifiable flexibility motif across all variants is the loop between B4 and B5 (residues 94-98) (**Figure 3A**). The THR crystal B-factors also locally increase in loops B2-H1(residues 48-57) and H1-B3 (residues 58-77). According to ML predictions, loops between secondary structure elements are clearly marked by an increase in protein flexibility, but there is no difference in flexibility between isoforms (**Figure 3B,D**). NMR RMSD is about one order of magnitude lower in the globular domain, compared to the IDD, and locally increases between secondary structure elements (**Figure 3C**). NMR J(0) analysis provides further insight into local flexibility. While NMR J(0) primarily categorises residues into “flexible” (IDD) and “rigid” (globular core), several residues within the globular domain deviate from this pattern. Notably, clear outliers appear at the C-terminal end of helix H1 and in the H1-B3 loop (**Figure 3E**). Classical triplicate MD and adaptive-sampling MD identify the same H1-B3 region as a flexibility hotspot, with reproducible RMSF patterns across simulations. Notably, RMSF has the same order of magnitude as NMR RMSD (**Figure 3F, H**). These methods also allow a more straightforward comparison between variants. While all three proteins exhibit similar fold and global flexibility patterns, SY and THR consistently display elevated fluctuations in the H1-B3 loop relative to WT. Hydrogen-deuterium RFU is roughly divided into 2 regions: residues 27-70 at RFU=0.2-0.5, residues 71-136 at RFU=0.5-0.8 with two dips in loop B3-B4 and B6 C-terminal strand (**Figure 3G**). Both regions display enhanced deuterium uptake on long timescales, indicating transient local unfolding or conformational exchange inaccessible to fast-timescale probes like NMR or MD.

Among the applied methods, HDX-MS, MD, and ML provide the clearest and most reproducible variant-to-variant differences, in part because they sample flexibility on complementary timescales and are unaffected by restraint-based biases from NMR RMSD or B-factors. In contrast, NMR J(0) robustly differentiates disordered regions from folded regions and highlights chemical exchange at the C-terminal of helix H1 and the H1-B3 loop. However, it is less sensitive to subtle mutational effects, except where chemical exchange takes place. Collectively, these methods paint a coherent picture of SAK flexibility: a universally flexible IDD; a relatively rigid globular core with well-defined flexible loops; and modest but consistent perturbations in local flexibility between variants, notably in the helix H1 and the loop H1-B3.

### Integrated analysis confirms differences in protein flexibility

To delineate the contributions of each technique to our understanding of SAK flexibility, we computed Spearman correlation coefficients on normalised flexibility data (**Figure 4A**). Overall, computational methods were correlated with high Spearman coefficients, while experimental methods were the most divergent. Crystallographic B-factors were the most correlated to computational methods, followed by HDX-MS data. Our explanation is that crystallographic B-factors can describe global protein vibration^61^, as opposed to local changes in protein flexibility, which can be simulated using our MD simulations, and can be therefore predicted by the employed ML models. HDX-MS data describes protein flexibility on the s-min timescale, whereas most other techniques describe protein flexibility at the ns timescale (with contributions from chemical exchange at the ms timescale for NMR J(0), which explains low Spearman correlations between HDX-MS and other techniques. NMR RMSD showed almost no correlation with any other technique, and NMR J(0) (which is a measure of rigidity, not flexibility) was weakly anti-correlated, especially to adaptive sampling MD, which samples a larger conformational space than classical MD. However, NMR J(0) measurements can be biased by chemical exchange at longer timescales, which can drastically affect interpretation. As a result, many residues for which NMR J(0) data was ambiguous were filtered out of the analysis. Finally, ML-based methods for predicting protein flexibility (Flexpert and BioEmu), which are in nature computationally inexpensive, correlate reasonably to MD, which is consistent with MD data being used for training both ML models^56,62^. Thus, they show promising potential in supplementing or replacing these expensive computational calculations on complementing the experimental view on protein flexibility.

**Figure 4.**
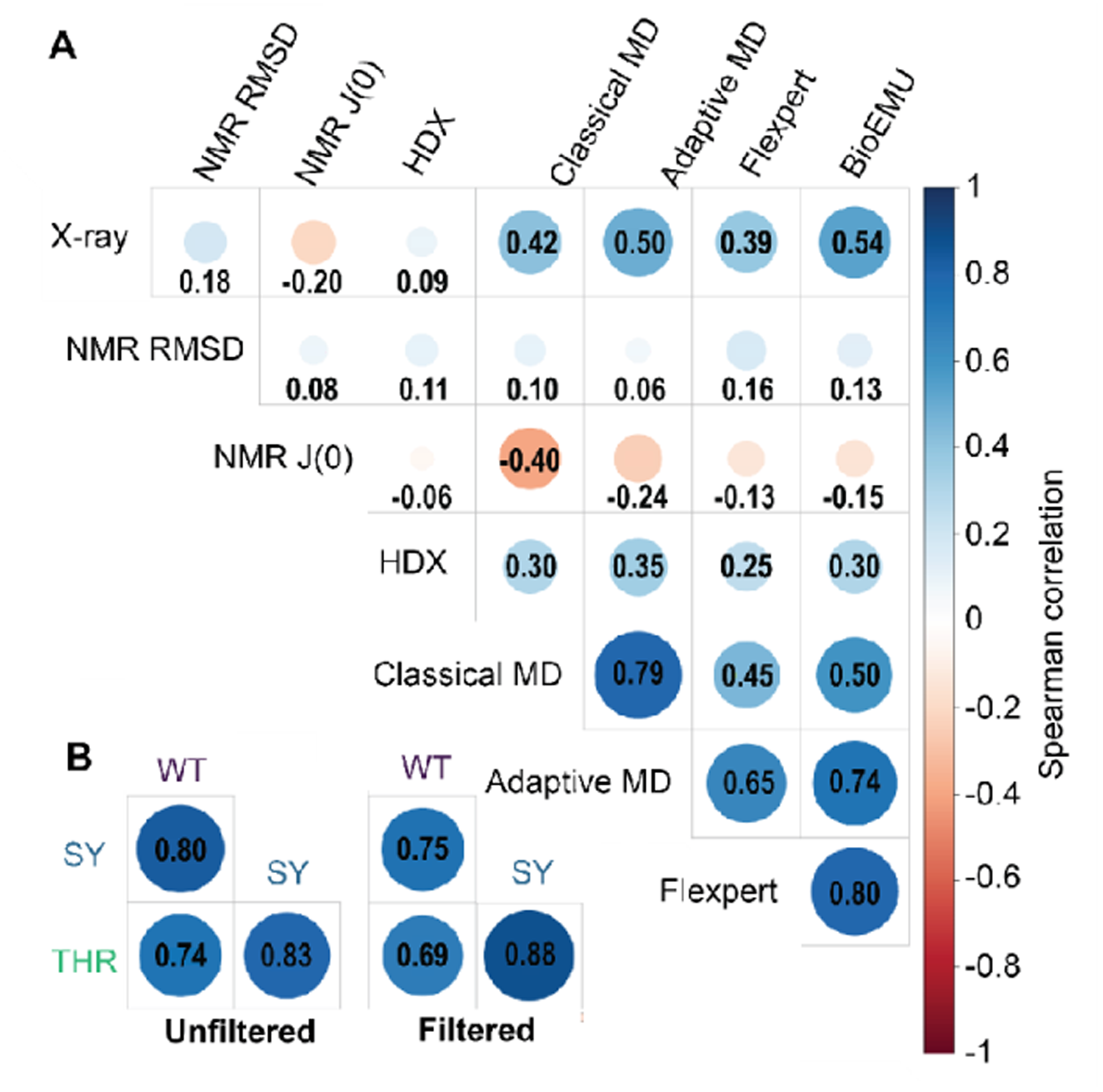
Comparing the flexibility of SAK variants. (A) Spearman correlations between flexibility datasets (concatenated) for all SAK variants. (B) Flexibility of SAK according to each dataset. (C) Spearman correlations between SAK variants (concatenated) for filtered (according to J(0)) residues (according to inability to obtain J(0) values).

Despite modest correlations in rankings of residue flexibility values, visualising protein flexibility according to each technique reveals consistent flexibility hotspots, notably the loop B1-B2, the loop H1-B3, and the loop B4-B5. (**Figure 4B**). Spearman correlations between SAK variants were inconclusive when integrating data at every position. Restricting our analysis to positions for which J(0) could not be computed (peak overlap, poor fit) revealed a clear difference in protein flexibility, where THR is the most divergent one compared to WT and SY (**Figure 4C**), suggesting that these positions (1-4, 6, 8, 9, 20, 23, 26, 28, 29, 35, 41, 42, 44, 46, 48, 51, 56, 57, 59, 60, 62, 63, 67, 70, 74, 76, 78, 79, 83, 89, 92, 104, 105, 108, 109, 114, 115, 128, 131, and 132) encompass individual amino-acids that present differential flexibility across variants.

## Discussion

Understanding how changes in local conformational and flexibility profiles contribute to staphylokinase function and immunogenicity is essential for rational design of efficient and safe thrombolytics. Although WT, SY, and THR share a conserved protein fold (**Figures 1B,2A**)^18,20^, our multimodal structural and biophysical characterization reveals differences in their oligomerization (**Figure 1A,C**), thermostability (**Figure 1D**), and flexibility (**Figures 3,4**). Most notably H1, B2, and adjacent loops distinguish the engineered low-immunogenic variants SY and THR from the WT.

The structural conservation among the three proteins is functionally critical, as all proteins exhibit similar plasminogen activation kinetics^63^. Thus, the multiple substitutions engineered into the SY and THR variants effectively reduce immunogenicity^12,13^ without compromising the fibrinolytic activity. NMR spectral density mapping, classical and enhanced sampling MD simulations, machine-learning predictions, and HDX-MS data indicate increased mobility around helix H1 in SY and THR compared to WT (**Figure 3**), where THR is a clear outlier (**Figure 4**), despite a preserved backbone topology. Since all three variants maintain similar plasminogen activation kinetics^63^, we can infer that: (i) increasing the flexibility of helix H1 does not significantly impact SAK-plasminogen binding and subsequent plasminogen activation, and (ii) substituted residues in SY and THR are not critical hotspots for plasminogen binding. Introduced substitutions and subsequent flexibility perturbations are localized in regions previously mapped as immunogenic hotspots^17^. Both SY and THR show modifications of the geometry of B2 strand (**Figure 2E**), which lies adjacent to the mapped antigenic sites, and exhibit locally elevated fluctuations in the H1-B3 loop region. Specifically, there is a local increase in NMR RMSD as well as in classical and adaptive MD RMSF, for the H1-B3 loop compared to WT, indicating an increase in flexibility in this key immunogenic region.

Most mutations in the low-immunogenic variants involve charge substitutions to alter the local charge distribution and potential hydrogen bonding, with the aim of reducing B-cell epitope recognition^12^. These mutations provide a clear molecular basis for the reduced immunogenicity and the altered flexibility profiles compared to WT and further support the correlation between flexibility perturbation and sequence modification. Taken together, our data indicates that mutations introduced in SY and THR produced only subtle structural changes but significant local changes in flexibility, notably on B2, H1, B3, and nearby loops.

The dimerization mode of SY observed in the crystal structure resembles that previously described by Chen *et al*. (PDB ID: 1C78)^64^ who proposed that such interactions could shield major antigenic epitopes. Our findings support this hypothesis that the SY variant forms dimers both in the crystal (**Figure 2C**) and in solution (**Figure 1A,C**), consistent with epitope masking as a mechanism for immunogenicity reduction. The putative dimerization interface of THR was shown to be primarily formed through [-strand interactions (**Figure 2D**), and has also been described previously by Chen et al.^64^ However, this study characterized this [-strand-[-strand interface as the weakest and least likely to occur under physiological conditions, suggesting that the THR dimer observed in crystal lattice is not biologically relevant. This supports the interpretation that the reduced immunogenicity in THR is driven primarily by flexibility changes or modification of the molecular surface. The NMR RMSD for THR was notably higher in the B3 region compared to the WT, suggesting that the substitutions destabilized the local backbone structure, leading to increased local flexibility in a region critical for B-cell recognition.

One of the challenges for protein engineers is the difficulty in accurately measuring and predicting protein flexibility, mainly because there is no consensus on the type of data to use as ground truth^65^ Our study highlights typical discrepancies between common data types used to describe protein flexibility. We used two machine learning-based methods relying on different approaches: BioEmu generates a conformational space from a single rigid structure^56^, while Flexpert was trained to predict on MD data RMSF of each residue from sequence^55^. Despite correctly assigning high flexibility to the N-terminal IDD and loops between secondary structure elements, both models failed to highlight any difference in variants (**Figure 3B,D**). In contrast, HDX-MS, NMR RMSD from structure ensembles, and classical / adaptive sampling MD data showed clear differences between variants, notably in B2, H1, B3, and the loops in between (**Figure 3C,F,G,H**). B-factors from X-ray crystallography were noisy and thus not very informative (**Figure 3A**). NMR J(0) was computed by reduced spectral density mapping from NMR relaxation data, and it clearly identified differences in rigidity at the ns timescale between the IDD and the globular domain of SAK but discriminated neither between secondary structure elements nor between variants (**Figure 3E**). Many J(0) values were detected in between H1 and B3, in accordance with previous observations by other techniques. Nonetheless, measurement of J(0) can be disrupted by peak overlap and slow / intermediate conformational exchange at the ms-s timescale, the latter causing peak broadening and hindering proper data fitting for residues involved^28^. This provided a useful filter to identify residues putatively undergoing slow / intermediate conformational exchange. Focusing on these residues, we showed that SY is the most similar to the WT in terms of flexibility (**Figure 4C**).

Overall, all computational predictions of flexibility were in good agreement with each other, with a clear bias for fast (ns time scale) protein flexibility. On the other hand, experimental techniques provided different picture of protein flexibility. We hypothesize this is due to the different timescales covered by each technique (**Figure 5**). The experimental technique showing the best correlation with computational ones was B-factors from X-ray crystallography, likely due to the fact that they are often used as a proxy to describe fast overall protein motion on the ps-ns timescale. This metric is however noisy and does not characterise local changes in flexibility^61^. HDX-MS, which relates deuterium uptake to protein flexibility at the second-minute timescale, is the second technique that correlates best with computational data. This would indicate that, there is a good correlation between fast and slow flexibility regimes in SAK. NMR RMSD from structure bundles was not correlated to any technique. RMSD is largely dependent on the number and quality of the distance restraints extracted from NOESY spectra, which themselves depend on the simulated annealing software used to compute the structure bundle, and the completeness of chemical shift assignment^66^. NMR J(0), a measure of rigidity at the ns timescale, is mildly anti-correlated to RMSF from classical and adaptive MD simulations, which measure flexibility at the ns timescale. Future attempts towards predicting SAK protein flexibility should focus on covering both fast and slow timescales, typically with a combination of ML predictors for fast ns regime, or MD simulations if higher precision is required to identify small differences, and HDX-MS at second-minute slow regime.

**Figure 5.**
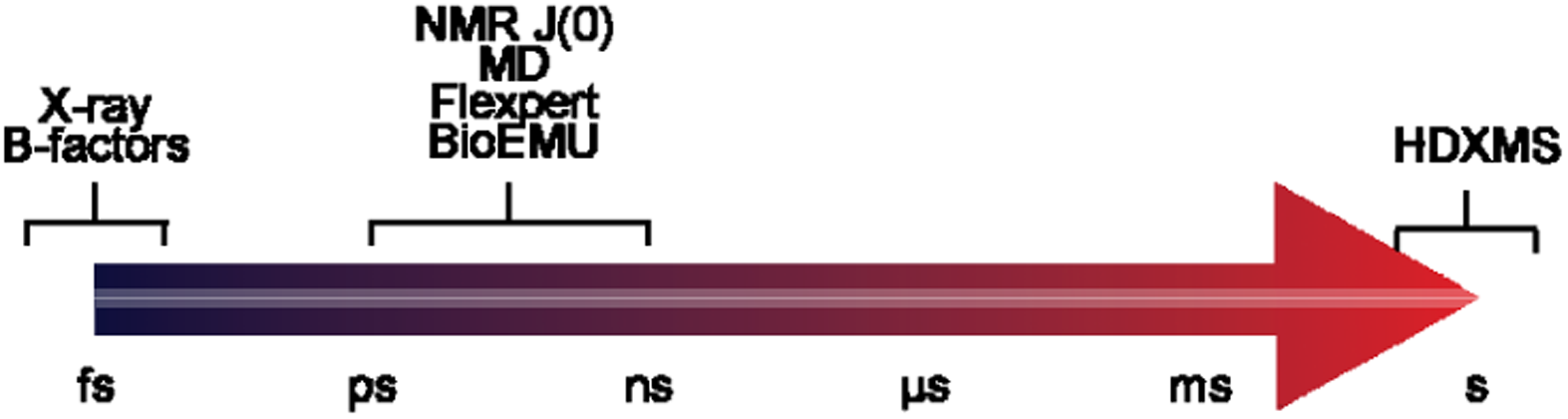
Summary of experimental and computational approaches used in this study mapped onto their characteristic time scales. Methods are arranged from femtosecond to second timescales, illustrating how agreement and disagreement between techniques arise from their intrinsic sensitivity to motions occurring on different timescales.

In summary, despite sharing a conserved fold and identical plasminogen activation kinetics, the low-immunogenic staphylokinase variants SY and THR exhibit altered oligomerization and pronounced local increases in flexibility, particularly in helix H1, strand B2, B3, and adjacent loops, compared to WT. Multimodal data collected using experimental (X-ray, NMR, HDX-MS) and theoretical (classical MD, accelerated MD, and ML) show that these flexibility perturbations localize to known immunogenic hotspots, providing a molecular basis for reduced immunogenicity without loss of function.

## Authors contributions

Anthony Legrand: isotopically-labelled protein production and purification, FIDA analysis,

NMR analysis, scientific illustration, conceptualization, manuscript writing

Linda Kasiarova: protein production and purification, biochemical and biophysical

characterizations, X-ray crystallography, scientific illustration, manuscript writing

Naina Verma: HDX-MS analysis, scientific illustration

Jan Mican: MD simulations, scientific illustration

Alan Strunga: protein production and purification for HDX-MS

Joan Planas-Iglesias: FlexPert, data analysis and interpretation, scientific illustration

Josef Kucera: HDX-MS analysis

Tomas Henek: HDX-MS analysis

Pavel Vanacek: HDX-MS analysis

Tim de Martines: Adaptive sampling MD simulations

Pavel Kohout : BioEmu data generation

Pavel Kaderavek: NMR analysis

Lukas Zidek: NMR analysis

Jiri Damborsky: conceptualization, funding acquisition, data interpretation

Stanislav Mazurenko: data interpretation, conceptualization

David Bednar: MD simulations

Lenka Hernychova: HDX-MS analysis, conceptualization

Zbynek Prokop: conceptualization, funding acquisition

Martin Marek: X-ray structure determinations, data interpretation, conceptualization, funding acquisition

All authors contributed to writing and review of the manuscript.

## Acknowledgements

The authors acknowledge the support of the RECETOX Research Infrastructure (No. LM2023069) and CZECRIN (No. LM2023049), funded by the Ministry of Education, Youth and Sports (MEYS) of the Czech Republic. This project was also supported by the European Union’s Horizon 2020 Research and Innovation Programme under grant agreements CETOCOEN Excellence (No. 857560) and CLARA (No. 101136607). Additional funding was provided through the ESIF-MEYS Johannes Amos Comenius Programme under the CLARA project (No. CZ.02.01.01/00/23_029/0008437), co-financed by the European Union and MEYS, and Czech Science Foundation (GX25-17329X and 25-18233M). The article reflects the author’s view, and the Agency and European Commission are not responsible for any use that may be made of the information it contains. Computational resources were provided by the e-INFRA CZ and ELIXIR-CZ (No. 90254 and LM2023055 MEYS). Anthony Legrand is supported by the Czech Ministry of Education, Youth and Sports under the OP JAK program (MSCAfellow5_MUNI, CZ.02.01.01/00/22_010/0003229). We acknowledge CF BIC and CF NMR of CIISB, Instruct-CZ Centre, supported by MEYS CR (LM2023042) and European Regional Development Fund-Project „Innovation of Czech Infrastructure for Integrative Structural Biology“ (No. CZ.02.01.01/00/23_015/0008175). The authors are thankful to ESRF synchrotron members for using their beamline facilities and help during data collection. LK, JM, and TM are supported by the scholarship Brno Ph.D. Talent.

## Data availability

Crystal structures of SAK SY and THR, and NMR structure bundles of WT, SY, and THR, were deposited to the PDB, under the following accession codes: 9IAU, 9IAV, 9RKK, 9RB5, 9RKL. NMR chemical shifts for WT, SY, and THR were deposited to the BMRB under the following accession codes: 34999, 34996, 35000. The mass spectrometry HDX data have been deposited to the Proteome Xchange (PX) Consortium^39^ via Proteomics Identifications (PRIDE)^40^ partner repository with the dataset identifier PXD072585; Username; Password: eLOcJZIiMDd9. The primary data from molecular dynamics simulations are available in Zenodo repository - https://zenodo.org/uploads/17853856

## References

1. Feigin, V. L. et al. Global, regional, and national burden of stroke and its risk factors, 1990–2019: a systematic analysis for the Global Burden of Disease Study 2019. Lancet Neurol. 20, 795–820 (2021).

2. Donkor, E. S. Stroke in the21stCentury: A Snapshot of the Burden, Epidemiology, and Quality of Life. Stroke Res. Treat. 2018, 1–10 (2018).

3. Feske, S. K. Ischemic Stroke. Am. J. Med. 134, 1457–1464 (2021).

4. Collen, D. & Lijnen, H. R. The Tissue-Type Plasminogen Activator Story. Arterioscler. Thromb. Vasc. Biol. 29, 1151–1155 (2009).

5. Tissue Plasminogen Activator for Acute Ischemic Stroke. N. Engl. J. Med. 333, 7 (1995).

6. Bivard, A. et al. Tenecteplase in ischemic stroke offers improved recanalization: Analysis of 2 trials. Neurology 89, 62–67 (2017).

7. Yaghi, S. et al. Treatment and Outcome of Hemorrhagic Transformation After Intravenous Alteplase in Acute Ischemic Stroke: A Scientific Statement for Healthcare Professionals From the American Heart Association/American Stroke Association. Stroke 48, (2017).

8. Toul, M., Nikitin, D., Marek, M., Damborsky, J. & Prokop, Z. Extended Mechanism of the Plasminogen Activator Staphylokinase Revealed by Global Kinetic Analysis: 1000-fold Higher Catalytic Activity than That of Clinically Used Alteplase. ACS Catal. 12, 3807–3814 (2022).

9. Nikitin, D. et al. Computer-aided engineering of staphylokinase toward enhanced affinity and selectivity for plasmin. Comput. Struct. Biotechnol. J. 20, 1366–1377 (2022).

10. Vanderschueren, S., Dens, J., Kerdsinchai, P. & Desmet, W. Randomized coronary patency trial of double- bolus recombinant staphylokinase versus front- Ioaded alteplase in acute myocardial infarction. 134, 7.

11. Vanderschueren, S. et al. A randomized trial of recombinant staphylokinase versus alteplase for coronary artery patency in acute myocardial infarction. The STAR Trial Group. Circulation 92, 2044–2049 (1995).

12. Laroche, Y. et al. Recombinant staphylokinase variants with reduced antigenicity due to elimination of B-lymphocyte epitopes. Blood 96, 1425–1432 (2000).

13. Pulicherla, K. K., Kumar, A., Gadupudi, G. S., Kotra, S. R. & Sambasiva Rao, K. R. S. In Vitro Characterization of a Multifunctional Staphylokinase Variant with Reduced Reocclusion, Produced from Salt Inducible E. coli GJ1158. BioMed Res. Int. 2013, 1–12 (2013).

14. Moussa, M. et al. Expression of recombinant staphylokinase in the methylotrophic yeast Hansenula polymorpha. BMC Biotechnol. 12, 96 (2012).

15. Collen, D. et al. Recombinant staphylokinase variants with altered immunoreactivity. I: Construction and characterization. Circulation 94, 197–206 (1996).

16. Collen, D. et al. Recombinant Staphylokinase Variants With Altered Immunoreactivity: III: Species Variability of Antibody Binding Patterns. Circulation 95, 455–462 (1997).

17. Jenné, S., Brepoels, K., Collen, D. & Jespers, L. High Resolution Mapping of the B Cell Epitopes of Staphylokinase in Humans Using Negative Selection of a Phage-Displayed Antigen Library. J. Immunol. 161, 3161–3168 (1998).

18. Rabijns, A., De Bondt, H. L. & De Ranter, C. Three-dimensional structure of staphylokinase, a plasminogen activator with therapeutic potential. Nat. Struct. Mol. Biol. 4, 357–360 (1997).

19. Ohlenschläger, O., Ramachandran, R., Gührs, K.-H., Schlott, B. & Brown, L. R. Nuclear Magnetic Resonance Solution Structure of the Plasminogen-Activator Protein Staphylokinase ,. Biochemistry 37, 10635–10642 (1998).

20. Parry, M. A. A. et al. The ternary microplasmin–staphylokinase–microplasmin complex is a proteinase–cofactor–substrate complex in action. Nat. Struct. Biol. 5, 917–923 (1998).

21. Lee, W., Rahimi, M., Lee, Y. & Chiu, A. POKY: a software suite for multidimensional NMR and 3D structure calculation of biomolecules. Bioinformatics 37, 3041–3042 (2021).

22. Klukowski, P., Riek, R. & Güntert, P. Rapid protein assignments and structures from raw NMR spectra with the deep learning technique ARTINA. Nat. Commun. 13, 6151 (2022).

23. Williams, C. J. et al. MolProbity: More and better reference data for improved all-atom structure validation. Protein Sci. 27, 293–315 (2018).

24. Kabsch, W. & Sander, C. Dictionary of protein secondary structure: Pattern recognition of hydrogen-bonded and geometrical features. Biopolymers 22, 2577–2637 (1983).

25. Ferrage, F. Protein Dynamics by 15N Nuclear Magnetic Relaxation. in Protein NMR Techniques (eds Shekhtman, A. & Burz, D. S.) vol. 831 141–163 (Humana Press, Totowa, NJ, 2012).

26. Raiford, D. S., Fisk, C. L. & Becker, E. D. Calibration of methanol and ethylene glycol nuclear magnetic resonance thermometers. Anal. Chem. 51, 2050–2051 (1979).

27. Emsley, L. & Bodenhausen, G. Gaussian pulse cascades: New analytical functions for rectangular selective inversion and in-phase excitation in NMR. Chem. Phys. Lett. 165, 469–476 (1990).

28. Kadeřávek, P. et al. Spectral density mapping protocols for analysis of molecular motions in disordered proteins. J. Biomol. NMR 58, 193–207 (2014).

29. Kabsch, W., XDS. Acta Crystallogr. D Biol. Crystallogr. 66, 125–132 (2010).

30. Evans, P. R. & Murshudov, G. N. How good are my data and what is the resolution? Acta Crystallogr. D Biol. Crystallogr. 69, 1204–1214 (2013).

31. Weichenberger, C. X. & Rupp, B. Ten years of probabilistic estimates of biocrystal solvent content: new insights via nonparametric kernel density estimate. Acta Crystallogr. D Biol. Crystallogr. 70, 1579–1588 (2014).

32. Adams, P. D. et al. The Phenix software for automated determination of macromolecular structures. Methods 55, 94–106 (2011).

33. Abramson, J. et al. Accurate structure prediction of biomolecular interactions with AlphaFold 3. Nature 630, 493–500 (2024).

34. Emsley, P., Lohkamp, B., Scott, W. G. & Cowtan, K. Features and development of Coot. Acta Crystallogr. D Biol. Crystallogr. 66, 486–501 (2010).

35. Moriarty, N. W., Grosse-Kunstleve, R. W. & Adams, P. D. electronic Ligand Builder and Optimization Workbench ( eLBOW ): a tool for ligand coordinate and restraint generation. Acta Crystallogr. D Biol. Crystallogr. 65, 1074–1080 (2009).

36. Peterle, D., Wales, T. E. & Engen, J. R. Simple and Fast Maximally Deuterated Control (maxD) Preparation for Hydrogen–Deuterium Exchange Mass Spectrometry Experiments. Anal. Chem. 94, 10142–10150 (2022).

37. Kavan, D. & Man, P. MSTools—Web based application for visualization and presentation of HXMS data. Int. J. Mass Spectrom. 302, 53–58 (2011).

38. Smit, J. H. et al. Probing Universal Protein Dynamics Using Hydrogen–Deuterium Exchange Mass Spectrometry-Derived Residue-Level Gibbs Free Energy. Anal. Chem. 93, 12840–12847 (2021).

39. Deutsch, E. W. et al. The ProteomeXchange consortium at 10 years: 2023 update. Nucleic Acids Res. 51, D1539–D1548 (2023).

40. Perez-Riverol, Y. et al. The PRIDE database resources in 2022: a hub for mass spectrometry-based proteomics evidences. Nucleic Acids Res. 50, D543–D552 (2022).

41. Anandakrishnan, R., Aguilar, B. & Onufriev, A. V. H++ 3.0: automating pK prediction and the preparation of biomolecular structures for atomistic molecular modeling and simulations. Nucleic Acids Res. 40, W537–W541 (2012).

42. Case, D. A., et al. AmberTools. J. Chem. Inf. Model. 63, 6183–6191 (2023).

43. Tian, C. et al. ff19SB: Amino-Acid-Specific Protein Backbone Parameters Trained against Quantum Mechanics Energy Surfaces in Solution. J. Chem. Theory Comput. 16, 528–552 (2020).

44. Izadi, S., Anandakrishnan, R. & Onufriev, A. V. Building Water Models: A Different Approach. J. Phys. Chem. Lett. 5, 3863–3871 (2014).

45. Darden, T., York, D. & Pedersen, L. Particle mesh Ewald: An N ⋅log( N ) method for Ewald sums in large systems. J. Chem. Phys. 98, 10089–10092 (1993).

46. Davidchack, R. L., Handel, R. & Tretyakov, M. V. Langevin thermostat for rigid body dynamics. J. Chem. Phys. 130, 234101 (2009).

47. Loncharich, R. J., Brooks, B. R. & Pastor, R. W. Langevin dynamics of peptides: The frictional dependence of isomerization rates of N -acetylalanyl- N 1-methylamide. Biopolymers 32, 523–535 (1992).

48. Berendsen, H. J. C., Postma, J. P. M., Van Gunsteren, W. F., DiNola, A. & Haak, J. R. Molecular dynamics with coupling to an external bath. J. Chem. Phys. 81, 3684–3690 (1984).

49. Roe, D. R. & Cheatham, T. E. PTRAJ and CPPTRAJ: Software for Processing and Analysis of Molecular Dynamics Trajectory Data. J. Chem. Theory Comput. 9, 3084–3095 (2013).

50. Waterhouse, A. et al. SWISS-MODEL: homology modelling of protein structures and complexes. Nucleic Acids Res. 46, W296–W303 (2018).

51. Doerr, S., Harvey, M. J., Noé, F. & De Fabritiis, G. HTMD: High-Throughput Molecular Dynamics for Molecular Discovery. J. Chem. Theory Comput. 12, 1845–1852 (2016).

52. Price, D. J. & Brooks, C. L. A modified TIP3P water potential for simulation with Ewald summation. J. Chem. Phys. 121, 10096–10103 (2004).

53. Maier, J. A. et al. ff14SB: Improving the Accuracy of Protein Side Chain and Backbone Parameters from ff99SB. J. Chem. Theory Comput. 11, 3696–3713 (2015).

54. KoubaPetr. Release Flexpert v1 · KoubaPetr/Flexpert. GitHub https://github.com/KoubaPetr/Flexpert/releases/tag/v1.

55. Kouba, P., et al. Learning to engineer protein flexibility. Preprint at 10.48550/arXiv.2412.18275 (2025).

56. Lewis, S. et al. Scalable emulation of protein equilibrium ensembles with generative deep learning. Science 389, eadv9817 (2025).

57. Taiyun. taiyun/corrplot. (2025).

58. R: The R Project for Statistical Computing. https://www.r-project.org/.

59. Krissinel, E. & Henrick, K. Detection of Protein Assemblies in Crystals. in Computational Life Sciences (eds R. Berthold, M., Glen, R. C., Diederichs, K., Kohlbacher, O. & Fischer, I.) vol. 3695 163–174 (Springer Berlin Heidelberg, Berlin, Heidelberg, 2005).

60. Gouet, P., Robert, X. & Courcelle, E. ESPript/ENDscript: extracting and rendering sequence and 3D information from atomic structures of proteins. Nucleic Acids Res. 31, 3320–3323 (2003).

61. De Sá Ribeiro, F. & Lima, L. M. T. R. Temperature-Resolved Crystallography Reveals Rigid-Body Dominance over Local Flexibility in B-Factors. ACS Omega 10, 38871–38881 (2025).

62. Vander Meersche, Y., Cretin, G., Gheeraert, A., Gelly, J.-C. & Galochkina, T. ATLAS: protein flexibility description from atomistic molecular dynamics simulations. Nucleic Acids Res. 52, D384–D392 (2024).

63. Strunga, A. Biochemical and Immunological Properties of Engineered Low-Immunogenic Staphylokinases for Next-Generation Thrombolytic Therapy.

64. Chen, Y. et al. Crystal structure of a staphylokinase variant: A model for reduced antigenicity. Eur. J. Biochem. 269, 705–711 (2002).

65. Guo, A. B., Akpinaroglu, D., Kelly, M. J. S. & Kortemme, T. Deep learning guided design of dynamic proteins. Preprint at 10.1101/2024.07.17.603962 (2024).

66. Fowler, N. J., Sljoka, A. & Williamson, M. P. A method for validating the accuracy of NMR protein structures. Nat. Commun. 11, 6321 (2020).

