## Supplementary Information File for "Investigating the Conformational Flexibility of Staphylokinase Across Multiple Time Scales"

**WT (SAK wild-type)**

SSSFDKGKYKKGDDASYFEPTGPYLMVNVTGVDSKGNELLSPHYVEFPIKPGTTLTKEKIEYYVEWALDATAYKEFRVVELDPSAKIEVTYYDKNKKKEETKSFPITEKGFVVPDLSEHIKNPGFNLITKVVIEKK

**SY (SAK SY155)**

SSSFDKGKYKKGDDASYFEPTGPYLMVNVTGVDSAGNELLSPHYVEFPIKPGTTLTKEKIEYYVQWALDATAYREFRVVELAPAAKIEVAYYDKNKKKDESKSFPITAAGFVVPDLSEHIKNPGFNLITTVVIERK

**THR (SAK THR174)**

SSSFDKGKYKKGDDASYFEPTGPYLMVNVTGVDSKGNELLSPHYVEFPIKPGTTLTKEKIEYYVDWALDATAYQEFEVVSLSPSAKIEVTYYDKNKKKEETKSFPITEKGFTVPDLSEHIKNPGFNLITYVVIRKK

**Figure S1.** Amino acid sequences of proteins used in this work.

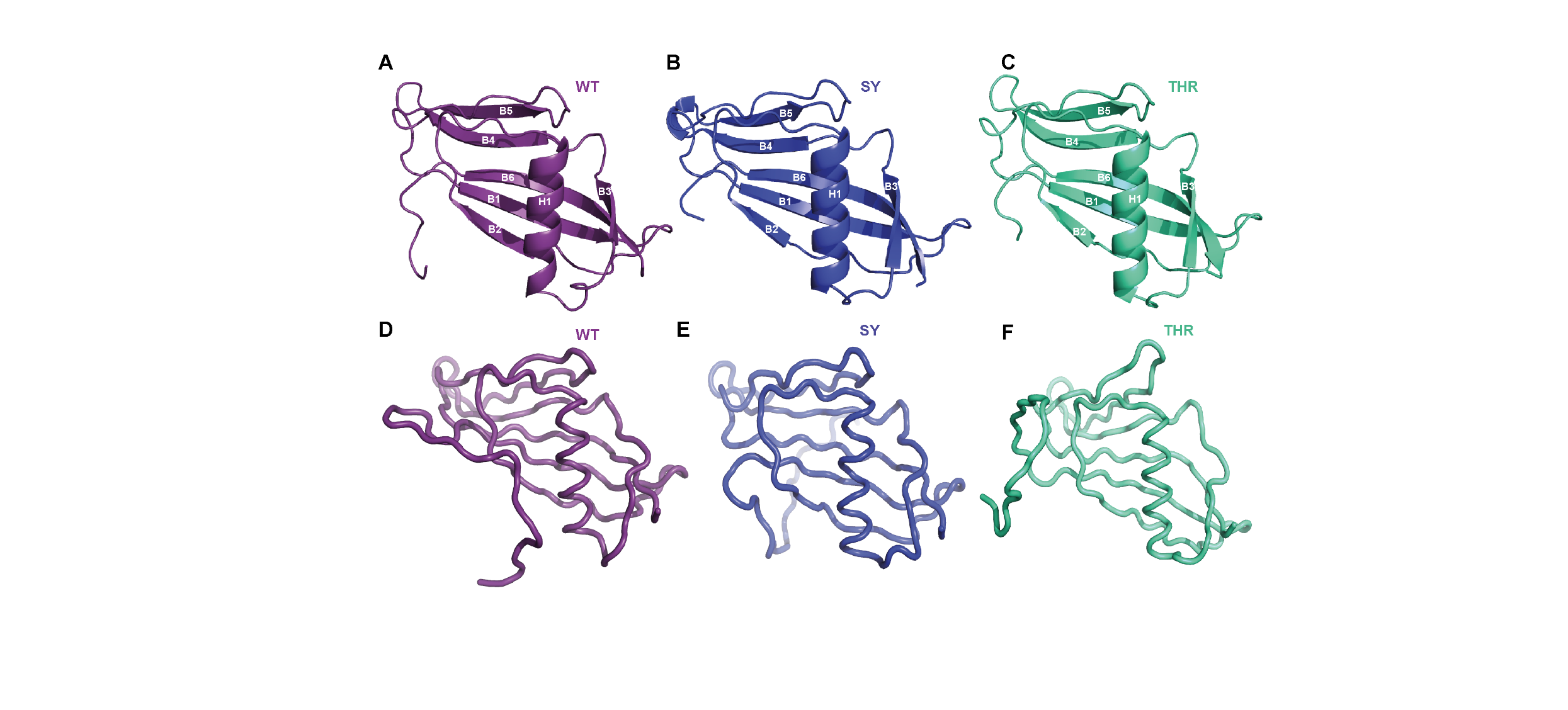

**Figure S2.** Crystal structures of (A) WT (PDB ID: 2SAK) (B) SY (C) THR and NMR structures of (D) WT (E) SY and (F) THR.

**Table S1.** Xray data collection statistics

| Data collection* | SAK-SY155 | SAK-THR174 |
| --- | --- | --- |
| Crystallization conditions | 0.18 M sodium citrate pH 4, 40% PEG 3350 | 0.18 M sodium citrate pH 4, 40% PEG 3350 |
| Wavelength (Å) | 0.9677 | 0.9677 |
| Resolution range | 44.31  - 2.09 (2.165  - 2.09) | 41.05  - 2.498 (2.588  - 2.498) |
| Space group | P 1 21 1 | P 61 2 2 |
| Unit cell parameters a, b, c (Å) | 46.007 79.477 79.876 | 51.283 51.283 322.902 |
| Unit cell parameters a, b, g (°) | 90 105.616 90 | 90 90 120 |
| Total reflections | 229844 (17366) | 180543 (18075) |
| Unique reflections | 30870 (2418) | 9657 (922) |
| Multiplicity | 7.4 (7.2) | 18.7 (19.6) |
| Completeness (%) | 93.94 (74.81) | 98.64 (91.01) |
| Mean I/sigma (I) | 10.98 (1.83) | 10.52 (0.67) |
| Wilson B-factor | 30.42 | 48.68 |
| R-merge | 0.1339 (1.027) | 0.2492 (2.366) |
| CC1/2 | 0.998 (0.542) | 0.997 (0.803) |
| Reflections used in refinement | 30870 (2418) | 9656 (840) |
| Reflections used for R-free | 1523 (129) | 468 (45) |
| R-work | 0.2331 (0.3018) | 0.2381 (0.4479) |
| R-free | 0.2741 (0.3305) | 0.2859 (0.6220) |
| Number of non-hydrogen atoms | 3739 | 2048 |
| Macromolecules | 3483 | 1970 |
| Ligands | 96 | 16 |
| Solvent | 160 | 62 |
| Protein residues | 439 | 243 |
| RMS (bonds) | 0.005 | 0.004 |
| RMS (angles) | 0.79 | 0.78 |
| Ramachandran favored (%) | 94.56 | 95.82 |
| Ramachandran allowed (%) | 3.55 | 2.93 |
| Ramachandran outliers (%) | 1.89 | 1.26 |
| Rotamer outliers (%) | 0.52 | 0.90 |
| Clash score | 8.78 | 3.79 |
| Average B-factor | 42.80 | 47.71 |
| macromolecules | 42.53 | 47.74 |
| ligands | 50.20 | 56.83 |
| solvent | 44.17 | 44.33 |
| PDB ID | 9IAU | 9IAV |

**Table S2.** NMR acquisition parameters.

| **Protein** | **^1^H Larmor** **frequency (MHz)** | **Experiment** | **Pulse sequence** | **Relaxation delay (s)** | **^1^H (90°) (μs)** | **^13^C (90°) (μs)** | **^15^N (90°) (μs)** | **Acqusition time (ms)** | **Time domain** | **Spectrum size** | **Window function** | **Shifted sinebell** |
| --- | --- | --- | --- | --- | --- | --- | --- | --- | --- | --- | --- | --- |
| WT | 950 | ^15^N-HSQC | hsqcetf3gp |  | 13.31 | 13.4 | 34 | 65.6 | 2048/256 | 2048/512 | QSINE | 2 |
|  |  | ^13^C-HSQC | hsqcgpph | 1 | 13.31 | 13.4 | 34 | 32.8 | 1024/256 | 1024/1204 | QSINE | 2 |
|  |  | CBCANH | cbcanhgpw3d | 1 | 13.31 | 13.4 | 34 | 77.8 | 2048/40/128 | 2048/128/256 | QSINE | 2 |
|  |  | CBCA(CO)NH | cbcacinhgpwg3d | 1 | 13.31 | 13.4 | 34 | 77.8 | 2048/40/128 | 2048/128/256 | QSINE | 2 |
|  |  | HNCA | hncagpwg3d | 1 | 13.31 | 13.4 | 34 | 77.8 | 2048/48/128 | 2048/128/256 | QSINE | 2 |
|  |  | HNCO | hncogpwg3d | 1 | 13.31 | 13.4 | 34 | 77.8 | 2048/40/128 | 2048/128/256 | QSINE | 2 |
|  |  | CC(CO)NH | hccconhgpwg3d3 | 1 | 13.31 | 13.4 | 34 | 65.6 | 2048/40/128 | 2048/128/256 | QSINE | 2 |
|  |  | H(CC)(CO)NH | hccconhgpwg3d2 | 1 | 13.31 | 13.4 | 34 | 65.6 | 2048/40/128 | 2048/128/256 | QSINE | 2 |
|  |  | ^15^N-NOESY | noesyhmqcf3gpph3d | 1 | 13.31 | 13.4 | 34 | 77.8 | 2048/40/128 | 2048/128/256 | QSINE | 2 |
|  |  | ^13^C-NOESY | noesyhsqcetgpsi3d | 1 | 13.31 | 13.4 | 34 | 77.8 | 2048/64/128 | 2048/128/256 | QSINE | 2 |
| SY | 700 |  |  |  |  |  |  |  |  |  |  |  |
|  |  | ^15^N-HSQC | hsqcetf3gp | 1 | 13.712 | 12 | 30 | 90 | 2048/256 | 2048/512 | QSINE | 3 |
|  |  | ^13^C-HSQC | hsqcgpph | 1 | 13.288 | 12 | 30 | 106 | 1024/256 | 1024/1204 | QSINE | 3 |
|  |  | CBCANH | cbcanhgpw3d | 1 | 13.288 | 12 | 30 | 106 | 2048/40/128 | 2048/128/256 | QSINE | 2 |
|  |  | CBCA(CO)NH | cbcacinhgpwg3d | 1 | 13.688 | 12 | 30 | 106 | 2048/40/128 | 2048/128/256 | QSINE | 2 |
|  |  | HNCA | hncagpwg3d | 1 | 13.688 | 12 | 30 | 106 | 2048/48/128 | 2048/128/256 | QSINE | 2 |
|  |  | HNCO | hncogpwg3d | 1 | 13.688 | 12 | 30 | 106 | 2048/40/128 | 2048/128/256 | QSINE | 2 |
|  |  | CC(CO)NH | hccconhgpwg3d | 1 | 13.288 | 12 | 30 | 90 | 2048/40/128 | 2048/128/256 | QSINE | 2 |
|  |  | HCCH-TOCSY | hcchdigp3d | 1 | 13.288 | 12 | 30 | 106 | 2048/58/128 | 2048/128/256 | QSINE | 2 |
|  |  | ^15^N-NOESY | noesyhmqcf3gpph3d | 1 | 13.688 | 12 | 30 | 106 | 2048/40/128 | 2048/128/256 | QSINE | 3 |
|  |  | ^13^C-NOESY | noesyhsqcetgpsi3d | 1 | 13.688 | 12 | 30 | 106 | 2048/64/128 | 2048/128/256 | QSINE | 3 |
| THR | 700 | ^15^N-HSQC | hsqcetf3gp | 1 | 12.831 | 12 | 30 | 90 | 2048/256 | 2048/512 | QSINE | 3 |
|  |  | ^13^C-HSQC | hsqcgpph | 1 | 12.831 | 12 | 30 | 71.7 | 1024/256 | 1024/1204 | QSINE | 3 |
|  |  | CBCANH | cbcanhgpw3d | 1 | 12.831 | 12 | 30 | 106 | 2048/40/128 | 2048/128/256 | QSINE | 2 |
|  |  | CBCA(CO)NH | cbcaconhgpwg3d | 1 | 12.831 | 12 | 30 | 106 | 2048/40/128 | 2048/128/256 | QSINE | 2 |
|  |  | HNCA | hncagpwg3d | 1 | 12.831 | 12 | 30 | 106 | 2048/48/128 | 2048/128/256 | QSINE | 2 |
|  |  | HNCO | hncogpwg3d | 1 | 12.831 | 12 | 30 | 106 | 2048/40/128 | 2048/128/256 | QSINE | 2 |
|  |  | CC(CO)NH | hccconhgpwg3d3 | 1 | 12.831 | 12 | 30 | 90 | 2048/40/128 | 2048/128/256 | QSINE | 2 |
|  |  | H(CC)(CO)NH | hccconhgpwg3d2 | 1 | 12.831 | 12 | 30 | 90 | 2048/40/128 | 2048/128/256 | QSINE | 2 |
|  |  | ^15^N-NOESY | noesyhmqcf3gpph3d | 1 | 12.831 | 12 | 30 | 106 | 2048/40/128 | 2048/128/256 | QSINE | 2 |
|  |  | ^13^C-NOESY | noesyhsqcetgpsi3d | 1 | 12.831 | 12 | 30 | 106 | 2048/64/128 | 2048/128/256 | QSINE | 2 |

**Table S3.** NMR structure bundle statistics and quality assessment.

| Resolution | WT | SY | THR |
| --- | --- | --- | --- |
| All atom | 0.93 ± 0.11 Å | 1.8 ± 0.2 Å | 1.9 ± 0.2 Å |
| All atom | 0.44 ± 0.49 Å | 1.1 ± 0.1 Å | 1.2 ± 0.7 Å |
| N | 0.16 ± 0.79 Å | 0.47 ± 0.13 Å | 0.80 ± 0.30 Å |
| CA | 0.17 ± 0.09 Å | 0.50 ± 0.14 Å | 0.84 ± 0.34 Å |
| CO | 0.17 ± 0.92 Å | 0.48 ± 0.14 Å | 0.81 ± 0.32 Å |
| CB | 0.20 ± 0.13 Å | 0.57 ± 0.15 Å | 0.95 ± 0.44 Å |
| First and last folded residues | 24-134 | 25-135 | 25-135 |
| Structural restraints |  |  |  |
| ^15^N-NOESY peaks | 3473 | 1662 | 1754 |
| ^13^C-NOESY peaks | 4298 | 1610 | 1366 |
| Distance restraints | 2167 | 1728 | 1659 |
| \|i-j\| = 0 | 535 | 436 | 473 |
| \|i-j\| = 1 | 632 | 585 | 549 |
| 1 < \|i-j\| < 5 | 290 | 232 | 191 |
| \|i-j\| >= 5 | 710 | 475 | 446 |
| Structure quality |  |  |  |
| ARTINA cycles (+ final CYANA structure calculation) | 7 | 7 | 7 |
| CYANA target function value | 20.90 ± 0.41 Å^2^ | 1.68 ± 0.12 Å^2^ | 2.14 ± 0.14 Å^2^ |
| Molprobity clashscore (percentile) | 14.12 ± 1.293693 (42^th^) | 0.3975 ± 0.3382 (99^th^) | 2.0430 ± 0.1069 (64^th^) |
| Ramachandran plot (%) |  |  |  |
| Favoured | 80 | 80.3 | 88.8 |
| Allowed | 15.6 | 19.7 | 9.4 |
| Outliers | 4.4 | 0 | 1.8 |
